# Neurons remap to represent memories in the human entorhinal cortex

**DOI:** 10.1101/433862

**Authors:** Salman Qasim, Jonathan Miller, Cory S. Inman, Robert E. Gross, Jon T. Willie, Bradley Lega, Jui-Jui Lin, Ashwini Sharan, Chengyuan Wu, Michael R. Sperling, Sameer A. Sheth, Guy M. McKhann, Elliot H. Smith, Catherine Schevon, Joel Stein, Joshua Jacobs

**Author notes:** Correspondence, 351 Engineering Terrace, Mail Code 8904, 1210 Amsterdam Avenue, New York, NY 10027.

## Abstract

The entorhinal cortex (EC) is known to play a key role in both memory and spatial navigation. Despite this overlap in spatial and mnemonic circuits, it is unknown how spatially responsive neurons contribute to our ability to represent and distinguish past experiences. Recording from medial temporal lobe (MTL) neurons in subjects performing cued recall of object–location memories in a virtual-reality environment, we identified “trace cells” in the EC that remap their spatial fields to locations subjects were cued to recall on each trial. In addition to shifting its firing field according to the memory cue, this neuronal activity exhibited a firing rate predictive of the cued memory’s content. Critically, this memory-specific neuronal activity re-emerged when subjects were cued for recall without entering the environment, indicating that trace-cell memory representations generalized beyond navigation. These findings suggest a general mechanism for memory retrieval via trace-cell activity and remapping in the EC.

## Introduction

The ability to organize our past experiences is a defining aspect of memory, and a crucial component of this is distinguishing between overlapping experiences for memory retrieval. For example, imagine that you have been asked to recommend things to do in a city you have visited frequently—the question elicits your memories of experiences from locations all around the city, and you can provide an answer without mixing up your memories for your different trips. While lesion studies have demonstrated that declarative memory processes depend on intact medial temporal lobe (MTL) structures, such as the hippocampus and entorhinal cortex (Scoville and Milner, 1957, Squire et al., 1993), it is not clear how the activity of neurons in these regions enables the brain to represent past experiences and distinguish between them. We examined the neural basis for dissociable representation of memories by examining spatial navigation, because the neural circuits and computations that support spatial navigation are thought to underlie memory processes more generally (Burgess, 2002, Buzsaki and Moser, 2013, O’Keefe and Nadel, 1978).

The discovery of place cells in the hippocampus (O’Keefe and Dostrovsky, 1971) and grid cells in the entorhinal cortex (Hafting et al., 2005), demonstrated spatial tuning for cells in regions that are essential to memory function (Squire et al., 1993). Spatially modulated neuronal activity has also provided evidence for a mechanism that might support memory representation and differentiation. Place fields “remap”—changing the location of their firing spatial firing fields in an environment in response to changes in sensory input for local cues or environmental structure (Leutgeb et al., 2005, Muller et al., 1987)—which demonstrates that place cells can differentiate between different spatial contexts by representing each context with a specific pattern of neural activity. Critically, researchers have proposed that remapping acts as a general mechanism for representing and differentiating non-spatial memories (Colgin et al., 2008). Recent work has supported this idea by showing that changes to an animal’s behavioral state, attention, or goal can induce also remapping (Dupret et al., 2010, Gauthier and Tank, 2018, Komorowski et al., 2009, Markus et al., 1995, Miller et al., 2013, Wood et al., 2000). This suggests remapping serves as a mechanism linking spatially responsive neurons to memory, such that cells remap in responses to changes in memory states.

We hypothesized that recalling different memories would elicit remapping of neuronal activity to the location of the specific memory being recalled. We further hypothesized that the memory-specific neural activity associated with remembered locations would be accessible even when subjects were not moving through the environment. In this way, we theorized that neurons in the MTL integrate the content and context of past experiences to represent and differentiate between memories—neuronal representations that persist beyond that environment for general memory retrieval.

To test this hypothesis, we recorded and analyzed the activity of single neurons from the MTL of human epilepsy patients as they performed a cued spatial-memory task in which they recalled the locations of cued objects while moving through a virtual environment. We observed a unique population of cells in the entorhinal cortex and cingulate, which we refer to as trace cells. Specifically, trace cells remap their activity to locations near the cued object–location memory, indicating that their neural activity related to the specific location relevant for the memory cued on each trial. Furthermore, as subjects moved through the cued object’s remembered location, the firing rate of entorhinal trace cell could decode the cued object for that trial, and this memory-specific neuronal activity was also present even when subjects were not moving through the environment. Trace cell activity in the entorhinal cortex thus illustrates a potential neural basis for the representation and differentiation of experiences for memory retrieval.

## Results

We recorded from 295 neurons in the entorhinal cortex, hippocampus, amygdala, and cingulate cortex of 19 neurosurgical patients performing an object–location memory task in a virtual, linear track environment (Fig. 1A). In this task, subjects were instructed to learn the locations of different objects along the track and then to recall the locations when the objects were removed. The task consisted of separate encoding trials and retrieval trials. Encoding and retrieval trials follow the same general structure and task instruction, except that objects are visible on the track during encoding trials, and are absent during retrieval trials. Each trial begins with an “cue period,” in which subjects view text instructions indicating the cued object for that trial. Following this is the “hold period,” during which subjects remain stationary at the entrance to the track for 4 seconds. Then, the “movement period” begins and the subject is moved automatically down the track. During encoding trials, the object remains visible on the track, allowing the subject to easily press a button as they approach the object’s location (Fig. 1B). During retrieval trials, the object is absent and subjects press a button at the location where they remember the cued object being present. Figure 1C shows that subjects performed this task accurately because they pressed the button within 2.8 virtual units of the correct location on average (7% of the track length).

**Figure 1:**
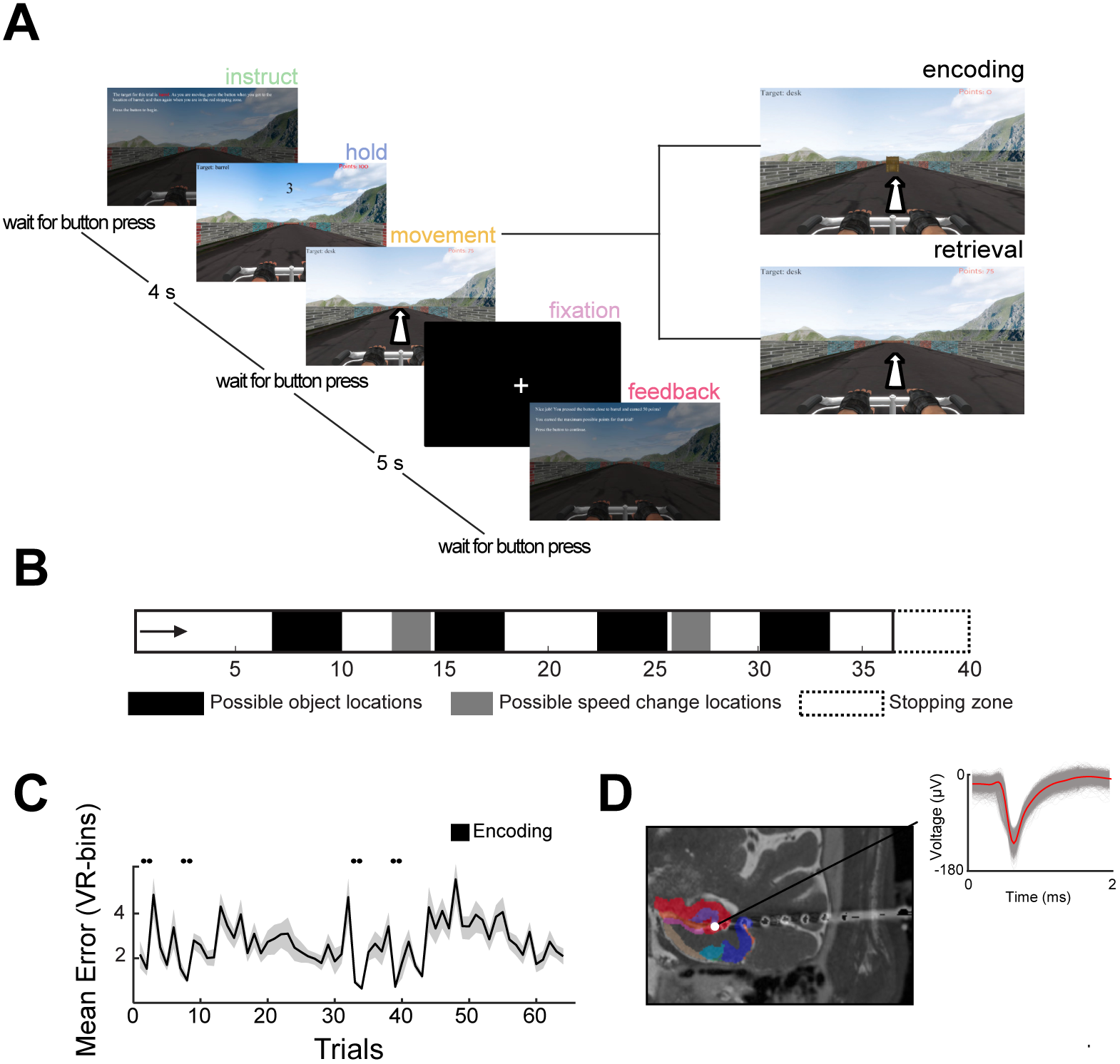
Task overview. A) Schematic of task design. Inset indicates that movement periods either consist of encoding or retrieval epochs. B) An overhead map of the environment. The arrow represents the starting point of each trial. C) Mean response error on retrieval trials across all sessions. Shading indicates SEM. D) MRI showing the electrode localization for a depth electrode in hippocampus. Shading indicates subregions: red, pink, or purple = hippocampus subregions, tan = entorhinal cortex, light or dark blue = perirhinal cortex. Inset shows example spike waveforms.

We examined the activity of each neuron in the task during retrieval trials by computing its firing rate as a function of the subject’s virtual location along the track. To assess the modulation of neuronal activity, we used a two-way repeated-measure ANOVA to identify neurons whose activity varied as a function of the subject’s location during retrieval trials, the retrieval cue, and their interaction. This analysis revealed two groups of neurons with distinct firing patterns. We found neurons with firing rates that varied as function of subject location alone (Fig. 2A), similar to conventional place cells (Ekstrom et al., 2003, O’Keefe and Dostrovsky, 1971). We also found a distinct cell type, which we call “trace cells,” that exhibited spatial firing fields that remapped to different locations along the track according to the retrieval cue on each trial (Fig. 2B, S2).

**Figure 2:**
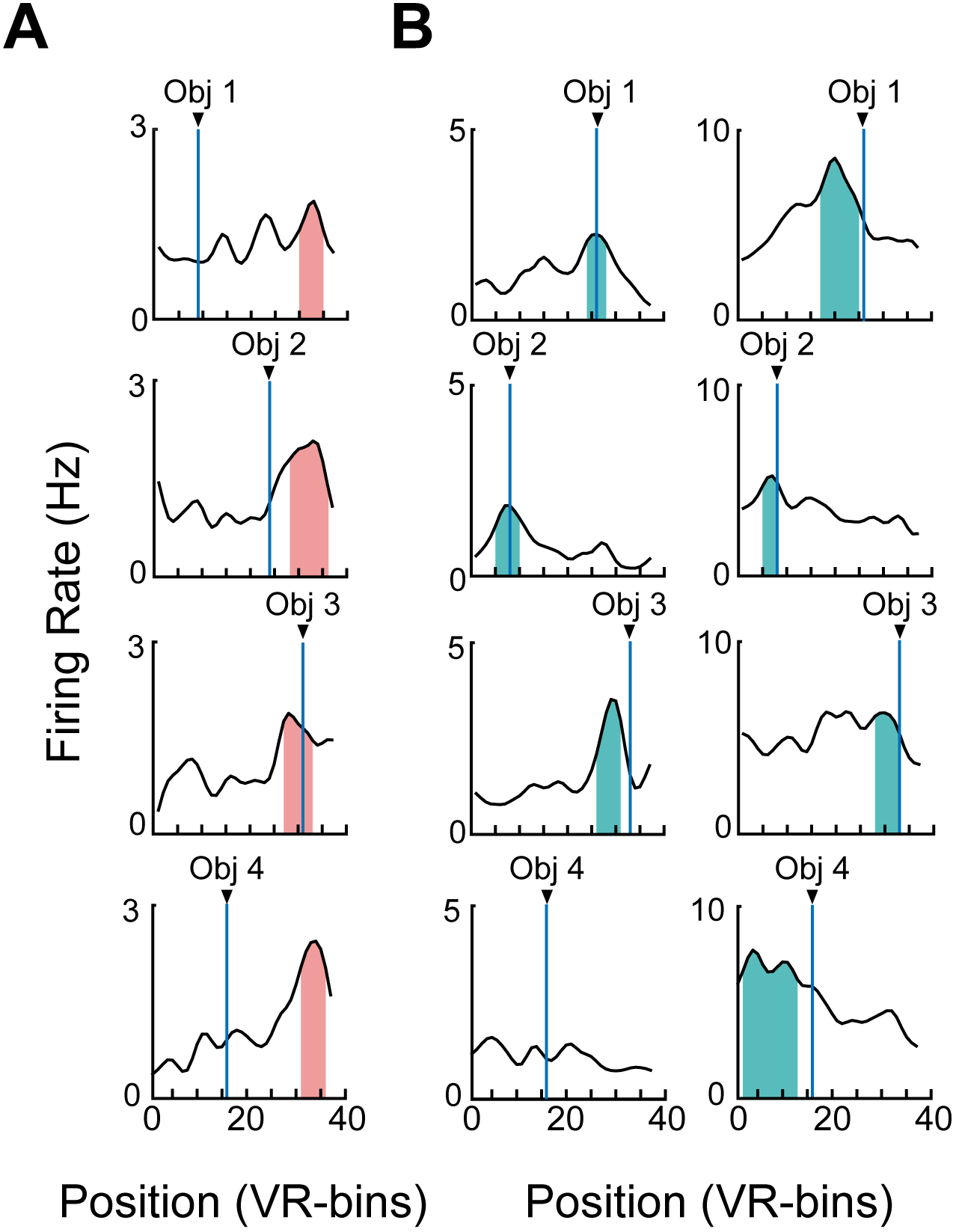
Examples of place and trace cells. A) The mean firing rate of an example hippocampal place cell. Individual plots show this cell’s activity for trial blocks with different cue objects. Vertical blue lines indicate object locations. Shading indicates statistically significant place fields (bins exceeding the 95th percentile of the shuffled firing rate distribution). B) The activity of two entorhinal trace cells. Note that in contrast to the place cells in panel *A*, these cells remap their firing fields depending on the object cue.

### Place cells activate in fixed locations, independent of memory retrieval demands

While subjects moved down the track, place cells activated in fixed locations of the environment (Fig. 2A, 3A). We defined place cells as those that showed a significant main effect of subject location on firing rate, and had at least one place field. We defined place fields by characterizing contiguous locations in which firing rate significantly exceeded a threshold measured with a permutation procedure (see *Methods*). A total of 16.9% of cells analyzed (50/295, *p* < 0.05, binomial test) showed this consistent spatial modulation of firing rate, and we classified them as place cells. A majority of spatial fields were smaller than 10% of the track length and none covered more than 40% of the track (Fig. 3C). We found significant numbers of place cells in the entorhinal cortex, hippocampus, and cingulate (Fig. 3D; binomial test, *p* < 0.05).

**Figure 3:**
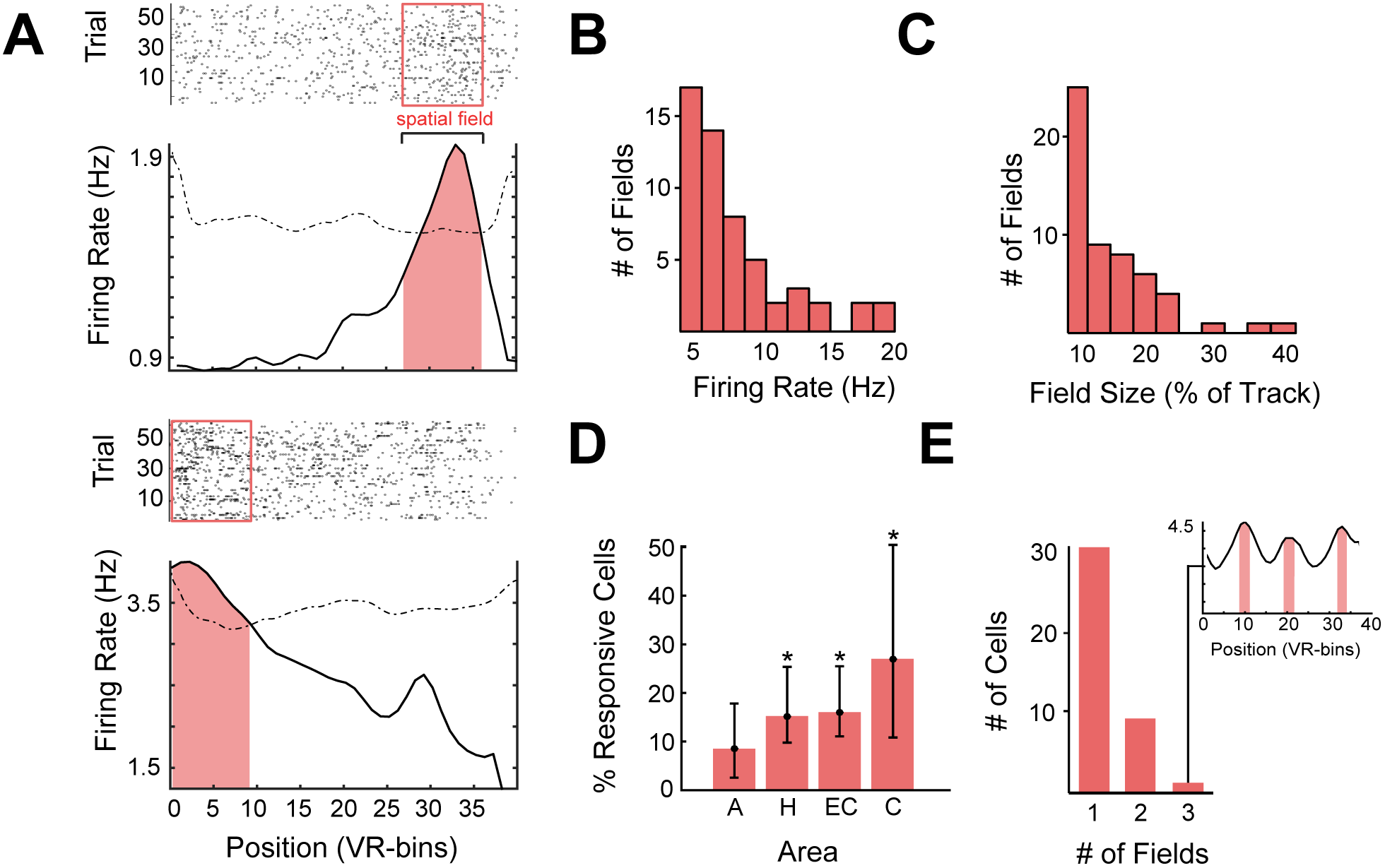
Place cell activity. A) Raster plot and mean firing rate for two representative place cells recorded from the hippocampus. Box and shading indicate location where the cell activity significantly increased. Dotted line represents the significance threshold, assessed with a shuffling procedure (see *Methods*). B) Distribution of mean firing rates among place fields. C) Distribution of field sizes as a percentage of the track. D) Proportion of place cells recorded in each brain area. A = amygdala, H = hippocampus, EC = entorhinal cortex, C = cingulate. Asterisks indicate location with a significant proportion of place cells (binomial test, *p* < 10−4). Bars indicate the 95% confidence interval from a binomial test. E)Number of responsive cells with more than one spatial field. Inset shows an example of multi–peak cell recorded from the cingulate.

Because 90% of the place cells continued to show this spatial coding even after accounting for potential effects time (MacDonald et al., 2011) or speed (Kropff et al., 2015), it indicates that a significant subset of responsive cells in the hippocampus, entorhinal cortex, and cingulate were modulated primarily by space rather than the memory demands of a trial.

#### Trace cells remap according to cued memory retrieval

In addition to place cells, we also observed trace cells whose firing fields remapped depending on the memory retrieval cue for each trial (Figs. 2B, S2). Figure 2B depicts two example cells recorded in the entorhinal cortex that showed spatially modulated activity. However, the particular location preference of each cell changed depending on the retrieval cue, or the specific object location that the subjects had been instructed to recall on each trial. Specifically, these cells significantly activated as subjects approached the cued object’s location, and then decreased afterwards. We defined trace cells as those that showed a significant interaction effect of the subject’s location and the retrieval cue on firing rate, and had at least one trace field. We characterized trace fields as the place field that a trace cell exhibited during the retrieval trials for a particular object location. We found significant numbers of trace cells (43/295; binomial test, *p* < 10^−11^), primarily in the entorhinal and cingulate cortices (Fig. 4A). We observed at least one trace cell in 15 of 19 subjects (Supp. Table 1); 12 of 19 subjects exhibited both place cells as well as trace cells.

**Figure 4:**
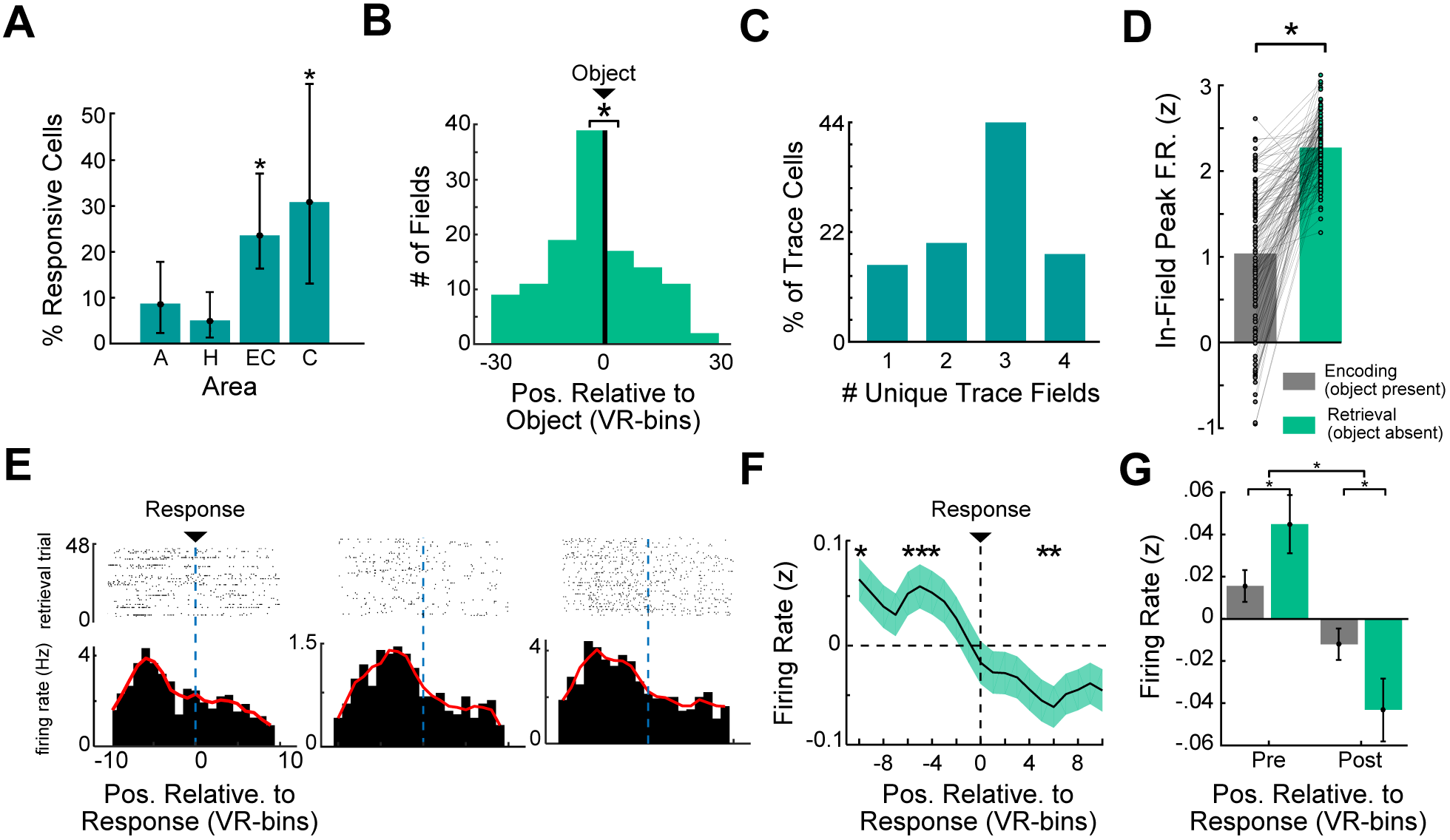
Trace-fields remap according to subjects’ memory for cued object locations. A) Distribution of trace cells across brain areas. Asterisks indicate significance at *p* < 10−5 (binomial test). B) Distribution of trace-field locations relative to object location (indicated by black line). Asterisk indicates the greater prevalance of trace fields immediately before versus after object location (χ^2^(1) = 10.4, *p* < 10^−3^). C) Distribution of the counts of unique trace fields exhibited by trace cells. D) Comparison of trace cell’s peak firing rate in field (z-scored) between encoding and retrieval trials (*t*(125) = 15.6, *p* < 10^−30^). E) Raster plot of spiking activity and corresponding PSTH for three representative entorhinal cortex trace cells, aligned relative to response location (indicated by blue dotted line). F) Mean firing rate (z-scored) of all trace cells aligned to response location. Shading indicates SEM. Asterisks indicate spatial bins that are significant from baseline (*p*’s< 0.05, one-sample *t* test, FDR-corrected). G) Pre-response and post-response firing rate (z-scored) compared between encoding and retrieval trials. Asterisks indicate significance from an ANOVA (interaction of pre-vs. post- and encoding vs. retrieval, *F*(1) = 5.79, *p* = 0.016).

The fact that trace cells remapped in response to changes in the memory retrieval cue seemed to demonstrate a possible mechanism whereby a single cell’s activity could maintain distinct representations of different memories. To test whether the particular nature of this remapping related to the specific object location that was recalled, we assessed where trace fields were most prominently located with respect to cued object locations during retrieval trials (when the object is no longer on the track). We found that trace fields were predominantly located preceding the cued object’s location (χ^2^(1) = 10.4, *p* < 10^−3^; Fig. 4B), which indicated to us that the activity of these cells could be driven by the memory for the object’s location. Critically, trace cells did not represent multiple remembered object locations simultaneously, instead switching between trace fields depending on the specific cued object (see Fig. 2B, S2). Trace cells did not always remap to the location of every cued object, with trace cells exhibiting anywhere from 1-4 trace fields throughout the session (Fig. 4C). These observations suggest that human trace cells remapped according to the retrieval cue—evidence that top-down memory retrieval demands influence remapping of trace-cell activity.

The findings described above left open the possibility that the activity of trace cells was driven by non-memory processes. Specifically, the activity of these cells might be explained by representations of object or goal locations (Deshmukh and Knierim, 2011, Gauthier and Tank, 2018, Hoydal et al., 2018, Sarel et al., 2017) or increases in visual attention related to object–scene associations (Moores et al., 2003). Each of these alternatives suggest that trace-cell activity would be conserved during encoding trials, which feature the same motor action, object, and goal location, but additionally provides visual cues in the form of the visible object on the track. We thus compared neural responses between retrieval and encoding trials because it allowed us to control for effects unrelated to memory retrieval. We examined trace cell firing rates as subjects passed through the center of each trace field during encoding versus retrieval trials and found that trace-cell firing activity was significantly greater during retrieval than encoding (*t*(125) = 15.5, *p* < 10^−30^; Fig. 4D). This significant increase in activity during retrieval suggests that trace cell activity reflected memory for object locations rather than visual responses to the object or it’s location.

These observations suggested that trace cells remap to cued object locations during memory retrieval, but did not directly link trace-cell activity to subjects’ memories for object locations. In order to assess whether trace-cell activity supports memory retrieval directly, we next assessed trace-cell activity relative to subjects’ response locations. Aligning trace-cell activity to subjects’s responses on retrieval trials, we found that trace cells showed increased firing in locations preceding the response location and then subsequently decreased their firing (Figs. 4E, S3A). This pattern was conserved when averaging over all trace cells Fig. 4F), though different cells tended to exhibit pre-response peaks at different distances to the response location (Supp. Fig. 3B). Trace cell activations thus correspond to subjects’ memory-driven responses for cued object locations, demonstrating that top-down memory demands drive the remapping of trace fields to cued object locations as shown in Figure. 4B.

An alternate explanation for these findings is that trace cells were activating in anticipation of subjects’ motor response (i.e., the button press). As before, we tested this possiblity by examining encoding trials when the motor demands identical to retrieval. During encoding trials we found significantly smaller changes in firing rates around the response location (ANOVA *F*(1) = 5.79, *p* = 0.016; Fig. 4G). This diminished effect in encoding trials indicates that trace-cell activity does not reflect anticipatory motor responses (see *Supp. Analyses* for additional controls).

### The firing rates of entorhinal trace cells distinguish between separate memories

In everyday life we often remotely recall events from outside of the environment in which they occurred. While our observation of trace cells above demonstrate that trace cell activity may scaffold distinct memories in their encoding environment, these findings do not show how they could be a useful mechanism for more generally dissociating memories without depending on movement through the encoding environment. We therefore asked if the same neuronal patterns associated with a particular memory emerge if subjects are cued for retrieval but do not move through the environment. To this end, we examined the activity of trace cells during the stationary hold period (Fig. 1A), when subjects are held at the beginning of the environment, which immediately followed cue presentation. Trace-cell firing rates during retrieval trials were significantly elevated during the hold period as opposed to all other periods of the task (Fig. 5A; ANOVA *F*(4) = 2.88, *p* = 0.02; FDR-corrected post-hoc *t*-tests *p* < 0.05), indicating that trace cells were possibly engaged by memory retrieval or maintenance related to the cued object during this period, even though subjects were not moving in the environment.

**Figure 5:**
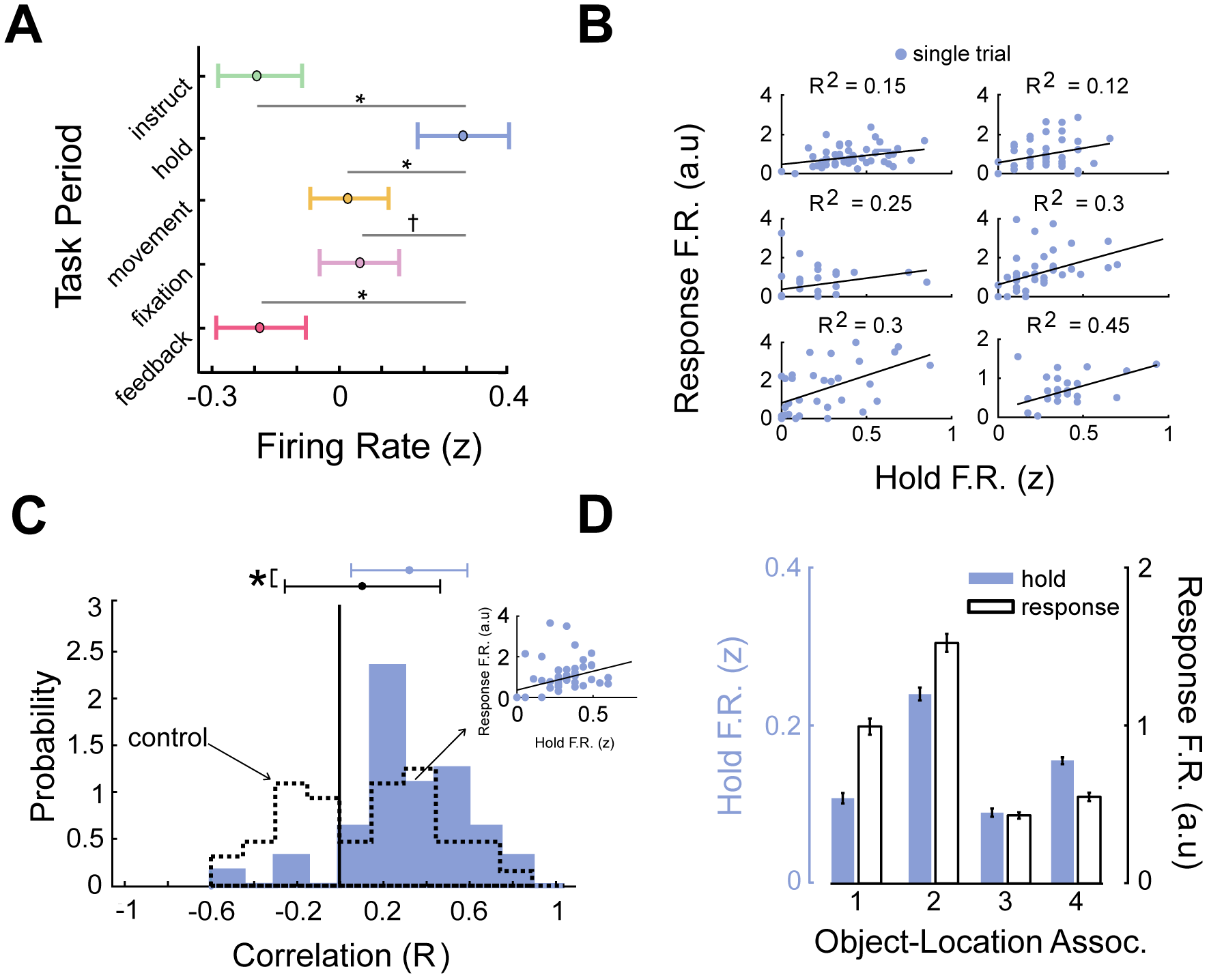
Trace-cell activity is correlated between the hold period and response period. A) Mean firing rate (z-scored) across all trace cells by task period. Asterisks indicate *p* < 0.05 (FDR corrected *t* tests); indicates *p* < 0.1. B) Relation between firing rates between hold- and response periods for six representative trace cells. Black line denotes the robust linear regression fit. C) Distribution of Pearson correlation coefficients for trace-cell firing rates between hold and response periods (mean = 0.31). Dotted line denotes control distribution (see *Methods* and *Supp. Analyses*). Asterisk indicates significant difference (*t*(42) = 6.5, *p* < 10^−9^). D) Mean normalized firing rate during hold and response periods for each object cue, for a representative entorhinal cortex trace cell.

If trace cells activate after cue presentation during the hold period, we hypothesized that this activity was related to the neural patterns associated with retrieval of cued object locations? If so, this would support the idea that trace-cell activity organizes memories in space but also generalizes beyond navigation to distinguish memories for retrieval. We therefore assessed if the trace-cell activity during the hold period correlated with the activity during the “response period” on the same trial, which is the period during movement when subjects responded to indicate the remembered object location. If trace cells were exhibiting a memory-specific rate code in response to the different retrieval cues, we reasoned that this level of neuronal activity should remain intact over both these periods. Consistent with our predictions, we found that trace-cell activity was positively correlated between these two periods within individual trials (Fig. 5B,C). This indicated that trace cells exhibited similar patterns of neural activity both during movement through the original encoding location and when subjects are held stationary at the entrance to the environment (e.g., Fig. 5D & S4).

To more directly demonstrate that the same patterns of neuronal activity in both the hold and response period consistently carried information about the memories being retrieved on each trial, we next used a cross-validated decoding framework to test if trace-cell activity was predictive of the content of object–location memories. This decoding analysis not only tested if activity in a single task period was able to reliably decode the cued object–location memory, but also whether a single shared neuronal representation of the current memory persisted across the hold and response periods, indicated by whether decoders trained in different settings reliably generalized to the response period neural activity. We trained decoders to use the normalized (*z*-scored) trace-cell firing rate from each task period (see Fig. 1A) to predict the identity of the cued object–location memory on each trial. We then tested each model’s decoding performance on neural activity from the response period (see *Methods*; Supp. Fig. 5). If we found significant classifier performance on this different test set, it would indicate that the same pattern of neural activity responded to specific object– location memories in a fashion that generalized across both periods. Decoding performance on the response period was significantly elevated for entorhinal cortex trace cells exclusively for the decoder trained on neural activity during the hold period (*p* < 0.01, binomial test; Fig. 6A). Critically, this indicates that the hold and response periods shared a common neural representation of the current cued memory. Non-entorhinal cortex trace cells exhibited chance-level decoding performance (Fig. 6B). This finding demonstrated that trace-cell activity in entorhinal cortex, specifically, represented spatial memories in a way that reliably dissociated different memories.

**Figure 6:**
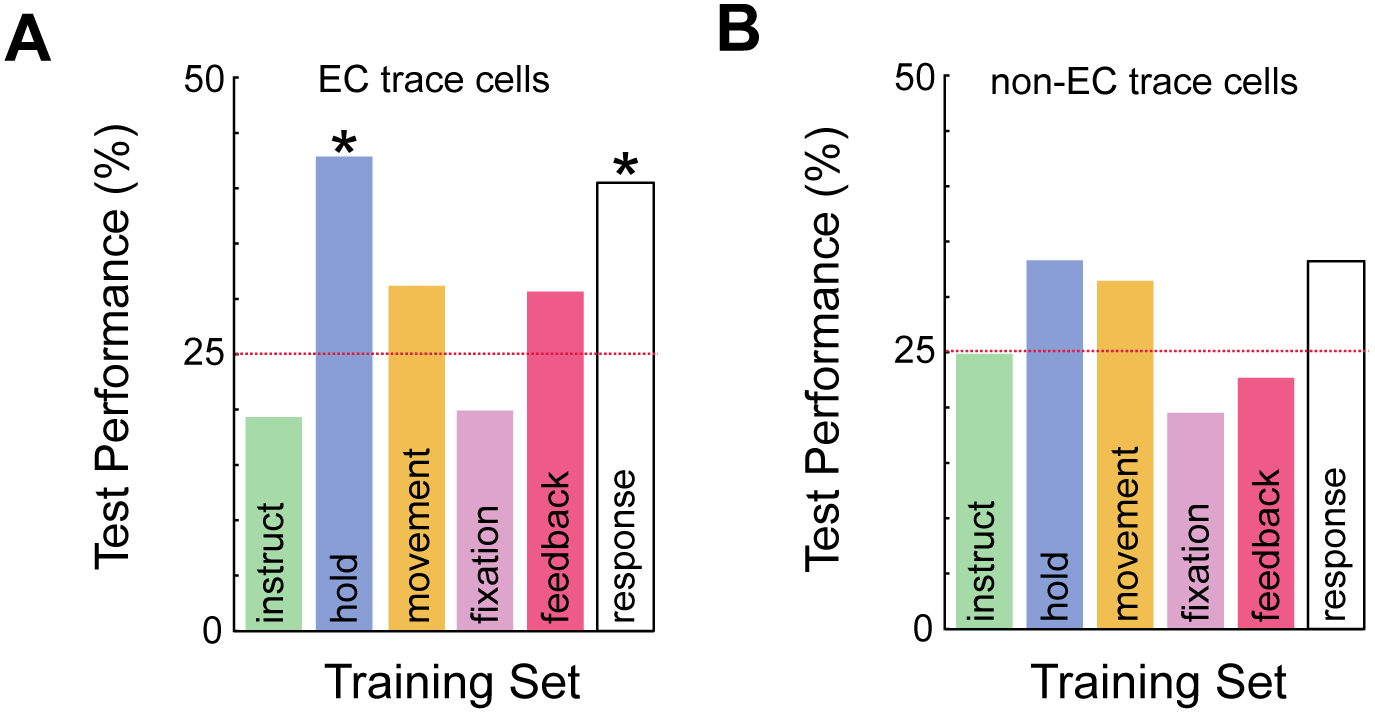
Trace-cell activity predicts memory content across hold period and response period. A) Results of memory-decoding analysis for entorhinal cortex trace cells. Individual bars distinguish models that were trained on different task periods, with all models tested on activity during the response period using trial-level cross validation. Red dotted line denotes chance (25%). Asterisks indicate above-chance decoding accuracy (binomial test, *p*’s< 0.02). B) Test performance for decoders based on non-entorhinal cortex trace-cell firing, with training on each of the task periods and testing on the response period.

## Discussion

A crucial aspect of human memory is our ability to differentiate and remember different past experiences. Here we show evidence that the activity of entorhinal trace cells represents and differentiates between memories from overlapping contexts. Critically, trace cells remapped their activity to represent locations that subjects had been cued to remember, illustrating a mechanism by which memory demands influence elements of the spatial map. These observations suggest that, as you move through an environment, trace cell activity plays a role in binding the objects and experiences you remember to the space in which they were present. Much of our memory recall does not occur while retracing our steps through the original environment, but outside of navigation through remembered locations. We found that trace cells were also active before subjects began moving through the environment on each trial, during the hold period following the presentation of the retrieval cue, suggesting that trace cells were not elicited purely by movement through space. The activity of entorhinal trace cells, specifically, was predictive of memory content during this stationary hold period, as well as when subjects moved through remembered object locations. That individual spatial memories were associated with a particular firing rate during both periods indicates that entorhinal trace cells are more generally involved in the dissociation and organization of memories beyond navigation. Below, we discuss how trace cells relate to previous single-cell findings in the hippocampus and entorhinal cortex relevant to space and memory, and help explain the role of the entorhinal cortex in the brain’s memory circuits.

Our findings share many features with the phenomenon of remapping as it is canonically described in rodent studies of spatial navigation. In these studies, changes to sensory cues relevant to the spatial environment induce changes in the location or rate of place field firing (Leutgeb et al., 2005, Muller and Kubie, 1987). In this way, place cells could potentially index different spatial maps for experiences in a particular spatial context. Given that different maps could conceivably re-activate during retrieval of those experiences, remapping was theorized to be a candidate mechanism for the indexing of different memories as well (Colgin et al., 2008). However, while past work has shown that behavioral or attentional state changes correspond to remapping in the hippocampus (Dupret et al., 2010), the relationship between remapping and memory has not been demonstrated. Our findings demonstrate neuronal activity that remaps as a function of changes to the memory subjects are cued to retrieve. This memory cue change occurs without any changes to local cues or the environment’s layout, indicating that memory processes alone can influence changes in spatial firing patterns in the human MTL.

This top-down influence on spatial firing bears similarity to work demonstrating that the locations of current goals can alter spatial firing patterns in the hippocampus (Eichenbaum et al., 1987, Gauthier and Tank, 2018, Komorowski et al., 2009, Sarel et al., 2017). In these studies, the activity of hippocampal cells in rodents and bats represented goal locations or vectors to goal locations. In contrast to goal or object cells in rodents or bats, trace cells in humans do not significantly activate when objects, the putative goal of the task, are visible in the environment. Rather, trace cells activate as subjects approach the remembered locations of these objects, when they are no longer visible on the track, indicating that trace cells are not tuned to visible goals. One possibility is that trace-cell activity represents a vector to remembered locations, similar to the activity of cells in the entorhinal cortex that represent vectors to objects in the environment (Hoydal et al., 2018).

The fact that entorhinal trace cells, in particular, were more generally predictive of memory content aligns with research in macaques showing that entorhinal cortex cells were active for maintenance and differentiation of item memories (Suzuki et al., 1997), and work in rodents describing the conjunctive representation of objects and space in the lateral entorhinal cortex (Deshmukh and Knierim, 2011). The trace cells we observed also resemble object-trace cells discovered in rodent entorhinal cortex (Knierim et al., 2014, Tsao et al., 2013) and cingulate (Weible et al., 2009). Object-trace cells in rodents are also active in locations where objects had previously been encountered, suggesting that the activity of these cells represent a non-specific, putative “memory trace” of the objects that the rodent had encountered in the environment, indicating “some object was here once.” In contrast, we show that trace-cell remapping in humans is driven by memory demands, leading to memory traces specific to the cued object location for memory retrieval.

The prevalence of trace cells in the entorhinal cortex helps us understand the importance of the entorhinal cortex in memory function. The entorhinal cortex is an early staging ground for attack by Alzheimer’s disease (Braak and Braak, 1991, Gomez-Isla et al., 1996, Khan et al., 2014, Masdeu et al., 2005). Recent evidence suggests that the spread of pathological tau through medial-temporal networks begins in the entorhinal cortex (Jacobs et al., 2018) and that entorhinal tau pathology is directly linked to memory decline in old age (Maass et al., 2018). It is unclear how, exactly, damage to the entorhinal cortex leads to a diseased memory state as few patients exhibit exclusively entorhinal lesions (Davachi, 2006,Olson and Newcombe, 2013). Our findings may help explain this issue. Given that Alzheimer’s disease disrupts the entorhinal cortex where trace cells are present, impaired trace-cell activity may play a role in the memory deficits that result from this disease. Furthermore, recording from trace cells during memory tasks may provide a direct way to assay entorhinal function during memory processes and prove useful in understanding the functional etiology of Alzheimer’s disease. Interestingly, while grid cells in the entorhinal cortex exhibit striking spatial responses (Hafting et al., 2005, Jacobs et al., 2013), recent imaging work shows reduced grid cell representations in patients at risk for Alzheimer’s (Kunz et al., 2015). Additionally, imaging studies have demonstrated that cognitive deficits related to both space and memory are associated with decline in healthy entorhinal function (Hirni et al., 2013, Olson et al., 2017, Reagh et al., 2018, Yeung et al., 2017). What relationship, if any, trace cells have to grid cells may be important to further understanding how memory influences the spatial map in humans.

In conclusion, our findings suggest that trace cells flexibly remap their spatial firing to distinguish individual memories. The fact that entorhinal trace-cell activity was predictive of memory content in both navigational and non-navigational settings shows that entorhinal cortex representations in memory extend more generally beyond navigation, which supports the notion that the entorhinal cortex is important for general relational and contextual memory representations (Behrens et al., 2018, Eichenbaum, 2014, Knierim et al., 2014, Lipton et al., 2007). Our findings may therefore enable new lines of electrophysiological investigation in various species of how neuronal representations of space and other domains are modulated by top-down demands from memory and other high-level processes.

## Methods

### Task

Nineteen patients with drug-resistant epilepsy performed 31 sessions of a spatial-memory task at their bedside with a laptop computer and handheld controller. In this virtual memory task, subjects are moved from the beginning to the end of a linear track on each trial. The track is 68 VR-units long, which roughly corresponds to 231 meters when converted using the height of the virtual avatar relative to the environment and track length. The ground was textured to mimic asphalt and the track was surrounded by stone walls (Fig. 1A). On each trial subjects are placed at the beginning of the track and shown text cues instructing them to press a button on the game controller when they reach the location of a specified object (“instruction period”). Immediately after receiving this cue, subjects press a button on a game controller to move to the “hold period,” in which they are held stationary at the entrance to the track for 4 seconds. Next, the “movement period” begins automatically, in which subjects are moved forward along the track. Subjects are moved passively for 56 of 64 trials and on other randomly selected trials control movements with a handheld controller (Supp. Fig. 1A)—we did not analyze the manual movement trials here. Individual trials consisted of either encoding or retrieval trials (see 1A). The first two times that subjects encounter a particular object are encoding trials, in which the object is visible during movement so the subjects can learn its location. On the subsequent retrieval trials, the object is invisible during movement and subjects are instructed to recall its location by pressing the controller button when they believe they are at the correct location. Subjects encode and retrieve a total of 4 unique object–location associations (16 trials of each) over the course of a session, with each object located at a different randomly selected location (Figure 1B). In addition to pressing a button to indicate their memory for the object location, subjects are told to press a button as they enter the “stopping zone” at the end of the track, which is visually delineated by a new floor coloring at the end of the track. Pressing the button in this region ends the movement period, and subjects are then shown a fixation cross for 5 seconds (“fixation period”). Finally, during the “feedback” period at the end of each trial, subjects receive points corresponding to how close they pressed the button to the correct location during movement. Only one object was ever present on the track at any given time. The task was split such that the retrieval cue for the first half of each session could correspond to objects 1 or 2, while retrieval cue for the second half could correspond to objects 3 or 4.

A distinctive feature of our task is that during movement periods subjects are moved subjects passively while their speed is automatically changed in a seemingly random fashion. These uncontrolled speed changes encourage subjects to attend continuously to their current location because they cannot accurately predict future positions by integrating their past velocity. Within each third of the track, subjects are moved at a constant speed, which is randomly chosen from the range of 2 to 12 VR units per second. The areas where speed changes occur is indicated in the schematic shown in Figure 1B. When speed changes occur, the speed varies gradually over the course of one second to avoid a jarring transition.

To measure task performance, we compute subject’s distance error (DE) on each trial, which is defined as the distance between the subject’s response location and the actual location of the object. We used a median split of each subject’s DE distribution to segment individual trials where performance was good versus bad.

#### Data recording

Epilepsy patients had Behnke–Fried microelectrodes surgically implanted in the course of clinical seizure mapping (Fried et al., 1999). Microwire implantation and data acquisition largely followed the procedures, approved by an Institutional Review Board (IRB), previously reported (Jacobs et al., 2013). Briefly, subjects undergoing treatment for drug-resistant epilepsy were implanted with clinical depth electrodes at four hospital sites: Emory University Hospital (Atlanta, GA), UT Southwestern Medical Center (Dallas, TX), Thomas Jefferson University Hospital (Philadelphia, PA), and Columbia University Medical Center (New York, NY). These electrodes feature 9 platinum–iridium microwires (40 μm) extending from the electrode tip and were implanted in target regions selected for clinical purposes. We recorded microwire data at 30 kHz using either the Cheetah (Neuralynx, Tucson, AZ) or NeuroPort (Blackrock Microsystems, Salt Lake City, UT) recording systems. We used Combinato (Niediek et al., 2016) for spike detection and sorting. We excluded neurons that had a mean firing rate below 0.2 Hz or above 15 Hz (potential interneurons). Manual sorting identified single-vs. multi-unit activity vs. noise on the basis of previously determined criteria (Valdez et al., 2013).

We determined the anatomic location of each implanted microwire electrode bundle using a combination of pre-implantation MRI and post-implantation CT scans. First, we performed automated whole brain and medial temporal lobe (Yushkevich et al., 2015) anatomic segmentation on T1-weighted (whole brain coverage, 3D acquisition, 1mm isotropic resolution) and T2-weighted (temporal lobe coverage, coronal turbo spin echo acquisition, 0.4 × 0.4 × 2 mm resolution) MRI. A post-implantation CT scan was then co-registered to the MRI scans and positions of electrode contacts and microwires were identified based on the source images and processed data (Jacobs et al., 2016).

### Identifying place cells and trace cells

To examine how neuronal activity varied with location in the virtual environment, we binned the virtual track into 40 bins, referred to as “VR–bins” (each VR–bin is equivalent to approximately 1.7 VR–units) enabling us to measure neuronal firing rates in this binned space. For each cell, we counted the spikes in each spatial bin and divided this quantity by the time spent in that bin to yield a firing rate estimate. We smoothed this firing rate estimate on the single-trial level using a Gaussian kernel with a width of 8 VR–bins. We excluded the bins in which subjects spent less than 100 ms over the course of the entire task. This excluded several of the bins in the stopping zone, because the movement period ended as soon as subjects pressed the button in the stopping zone. We normalized firing rate for all analyses comparing spiking across different task periods or trial types, such that a z-score of 0 represented a cell’s mean firing rate across all periods of the task.

We used a 2-way repeated-measure ANOVA to examine the effects of subject location (1–40 VR-bins), object cue (1, 2, 3, 4), and their interaction, on the binned firing rate of each cell. We defined place cells as those that showed a consistent and significant main effect of location on firing rate via the ANOVA, and that also had a place field greater than 5% the size of the track. Additionally, we performed an ANCOVA to confirm the main effect of position in the ANOVA, with position serving as a main factor while speed and time were covariates (Robitsek et al., 2013). We only considered a neuron to be a place cell if its firing was significantly modulated by place even after factoring time and speed in as covariates in the ANCOVA. We defined place fields as regions at least 5% the size of the track where the firing rate was significantly elevated (Ekstrom et al., 2003). To robustly determine statistical significance, we used permutation testing to build empirical estimates of the null distributions from which the test statistics could be drawn to determine significance from the real data. This shuffled distribution was created by circularly shifting the firing rate estimates 500 times and re-analyzing the data. Six cells showed a main effect of object cue on firing rate. These cells were excluded from analyses.

We defined trace cells as those cells whose firing rate showed an interaction between subject location and object cue in the ANOVA. Trace fields in trace cells were determined via the same method as for place cells, using a post-hoc test to identify firing fields that were specific to individual object–location associations. A trace field for a particular object cue was considered unique if the peak location did not overlap with that of any other trace field for that cell (Fig. 1D).

### Decoding analysis

We used a multivariate decoding framework to test whether trace-cell activity reflected information about the content of each object–location memory across different retrieval contexts. This framework is schematized in Supplementary Figure 5. To assess decoding performance, we pooled the trace cells recorded across all patients and sessions and constructed two pseudopopulations: entorhinal trace cells and non-entorhinal trace cells. Pseudopopulation decoding has been used to describe the common neural dynamics of functionally similar subsets of cells without the inherent noise correlations shared by neurons recorded in the same session (Kamiński et al., 2017).

The purpose of this decoding analysis was to ascertain whether a group of neurons provided a representation of the contents of memory that was similar in form across across separate contexts. For this decoding, we used a k-nearest neighbors (kNN) algorithm using a one-vs.-all paradigm for multi–class decoding of the identity of the remembered item from the recorded neuronal activity. Firing rates were binned by task period and normalized. On each trial we computed the “response period” firing rate by normalizing the activity in the 10 VR–bins preceding the response by the 10 VR–bins following the response (Supp Fig. 5). This normalization procedure captured both the pre-response increase and post-response decrease in firing rate described in the results. We used a similar method to compute a matched “control period” utilized in Figure 6C, using the 20 VR–bins immediately following the end of the response period. This ensured that the control period was of equal length to the response period, and that the neural activity during this control period did not overlap with the neural activity during the response period.

We trained all the different task period decoders on the firing rate during a particular period of the task and tested on the response period neural activity. Additionally, we trained and tested one decoder with the response period firing rate - this decoder was trained using leave-one-out cross validation to assess performance (Supp Fig. 5). We assessed significant decoding accuracy using a binomial test. Chance-level decoding accuracy was at 25%, given the equal presentation of the 4 different objects.

## Acknowledgements

We are grateful to the patients for participating in our study. This work was supported by NIH grants R01 MH104606, and S10 OD018211, and NSF Graduate Research Fellowship DGE 16-44869. We thank Andrew Watrous, Melina Tsitsiklis, and Ida Momennejad for helpful comments.

## Author contributions statement

J.J. conceived the experiment, R.B.G., J.T.W., B.L., A.S., C.W., S.A.S., G.M. performed surgical procedures, S.Q., J.M., M.S., C.S., E.S., J.J., C.I., performed data collection and recording, J.S. processed neuroimaging data, S.Q. analyzed the data, S.Q. and J.J. wrote the manuscript.

## Supplementary Analyses

### Control: Binning firing rate by space

In order to assess the spatial binning on our results, we calculated the results of our main analyses using 30 and 50 equal sized spatial bins, rather than 40 bins as in our main analyses. The number of place cells and trace cells identified by the ANOVA did not vary significantly as a function of the number of bins (results remained within 95 % confidence interval of binomial test determining significant proportion of place and trace cells). This indicates that our primary results are not determined by the spatial scales of the bins used for data analysis.

### Control: Electrodes in epileptic regions

The subject cohort examined in this study has drug-resistent epilepsy. Prior research has supported past work in epileptic cohorts through use of scalp EEG or fMRI (Lachaux et al., 2003). Still, it is important to consider whether electrophysiology research in the epileptic brain is reflective of healthy brain. Approximately 31% of the single-units we analyzed were recorded on microwires localized to clinically determined ictal onset zones. To more rigorously control for any confounding effect of epileptic activity, we re-ran all analyses excluding all neurons recorded from these clinically defined ictal onset zones. Our main findings remained unchanged with respect to the proportion of place cells and trace cells, and their properties. Further, this data exclusion did not change any results with respect to trace-cell activity or decoding.

### Control: Independence of multiple sessions by a single patient

Several patients contributed multiple sessions of the task, with each session analyzed independently. However, in order to ensure that patients contributing multiple sessions to this study were not confounding our results (Supp. Table 1), we ran control analyses utilizing only the first session recorded from each patient. This controlled for any confounding effect of multiple sessions. Our main findings remained unchanged with respect to the proportion of place cells and trace cells, and their properties. The results presented here thus utilize all the data.

### Trace cell activity follows subjective memory judgment

We sought to understand whether trace-cell activity followed the participants’ subjective memory of the object-location regardless of whether that memory was correct or incorrect. We tested this by splitting the retrieval task data into ”good” and ”bad” memory trials utilizing a median split within each subject. Both good and bad retrieval trials showed the same pattern of trace-cell activity with respect to response locations (pre, post paired t-test *t*(978) = *−*0.43*, p* = 0.66 *t*(978) = 1.12*, p* = 0.26; Supplemental Fig. 3C) — firing elevated before subjects’ response and decreased after, regardless of whether the trial was from the best half or worst half of the subjects’ performance. Given that we determined trace-cell activity was not an effect of the button press action itself, this suggested that the trace cells track a person’s subjective memory of the object-location, whereas if these cells were involved in context reinstatement we likely would have observed less activity during bad memory trials.

### Control Analysis: Trace cells do not encode time to button press

One alternative explanation of trace-cell activity is that it reflects a fixed anticipatory signal for the subjects’ motor action, the button press. Given that every trial featured random speed changes, our task controlled for consistent effects of time. This inherently meant that trace cells activating at consistent locations relative to subjects’ response were not responding at consistent times relative to that response, as time and location were dissociated across trials. To illustrate this, we assessed trace cell firing as a function of time relative to subjects’ response. We analyzed the activity of the trace cells time-locked to button press, rather than aligning trace cell activity by spatial bin/distance to button press as in Figure 2F. Anticipatory motor responses are thought to occur within 1-second preceding the relevant event (Mauritz and Wise, 1986), so we analyzed trace-cell firing in a 3-second window surrounding the response. Supplementary Figures 3D,E shows that trace cells did not show any consistent effect of time, as opposed to Figure 2D, in which trace cells exhibit clear preference for particular spatial positions that preceded retrieval. These results provided further evidence that the activity of trace cells reflected spatial activations at or near the remembered positions of cued objects, rather than simply firing at a fixed time preceding button press.

### Control: Trace cell hold–response period correlation does not result from temporal auto-correlation

Given that the response period activity was calculated by normalizing the pre-response firing rates by the post-response firing rates, the results in Fig. 4B,C already control for the effects of temporal autocorrelation (i.e., the hold period firing rate predicts the firing rate for the rest of the trial). To further ensure that the correlation we observed between the hold period firing rate and the response period firing (see Fig. 4B,C) was not the result of such a confound, we computed the correlation between the hold period firing rate and a ”control period”. Control period activity was computed using the length of the track following the response period, thus ensuring it used the neural activity in the regions of the track that did not overlap with the response period. This control period firing rate was computed identically to the response period—the mean firing rate of the first 10 VR–bins of the control period were normalized by the mean firing rate of the last 10 VR–bins. We then computed the correlation between the hold and control period firing. The null distribution of correlation coefficients assessed in this way is depicted by the dotted line in Fig. 5C.

**Supplementary Figure 1:**
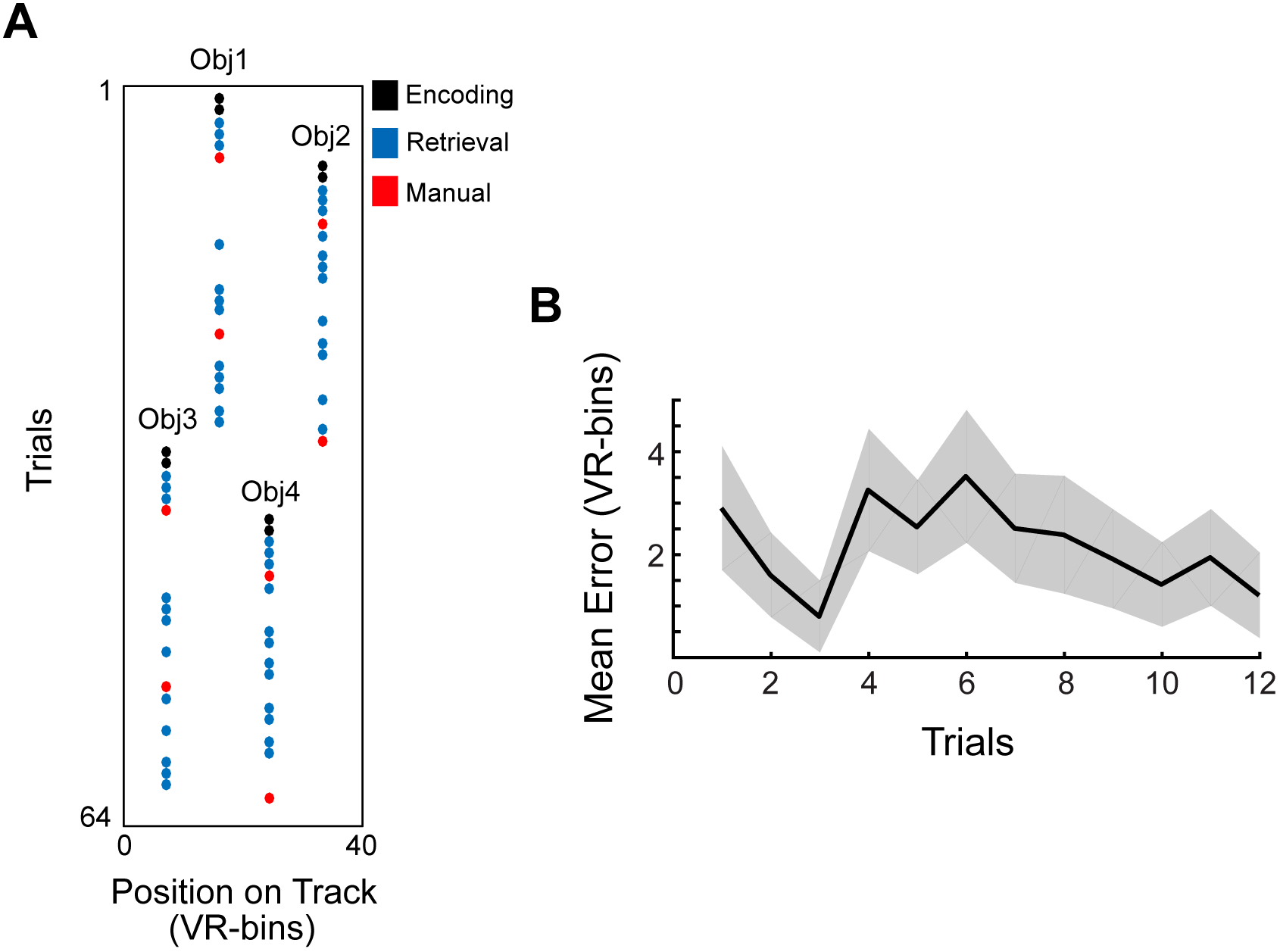
Trial structure and average response error: A) Schematic depicting one example of the trial structure during the task. The first two trials for each object cue were encoding trials (black), after which subjects had retrieval trials (passive movement, blue, manual movement, red). The cued object switched between objects 1 and 2 during the first half of the task, or objects 3 and 4 for the second half. Across sessions, the trials for each cue was random. B) Response error averaged across the 12 retrieval trials (blue dots seen in panel *A*) comprising each object–cue block. Shading indicates SEM. Note that subjects learned the object–location association quickly (first two trials), only to decrease in performance upon the introduction of another object location to hold in memory.

**Supplementary Figure 2:**
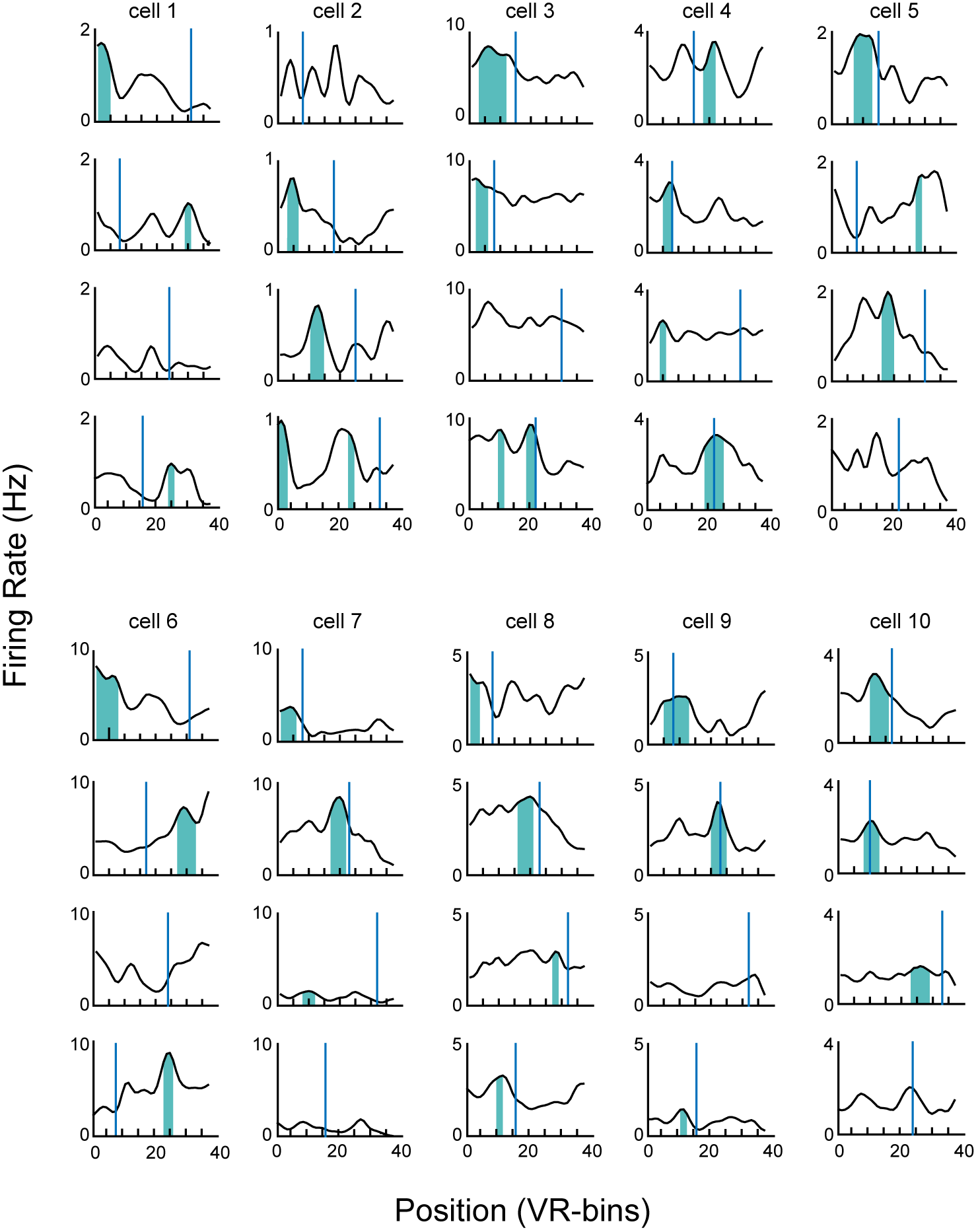
Examples of trace cell activity relative to object locations. Plots illustrate mean firing rates of trace cells for each object cue, as in Fig. 1F. Vertical blue lines indicate object locations. Shading indicates significant trace fields, as identified with a shuffling procedure (see *Methods*). Regions: cell 1 = amygdala, cells 2–3 = cingulate, cells 3–9 = EC, cell 10 = hippocampus.

**Supplementary Figure 3:**
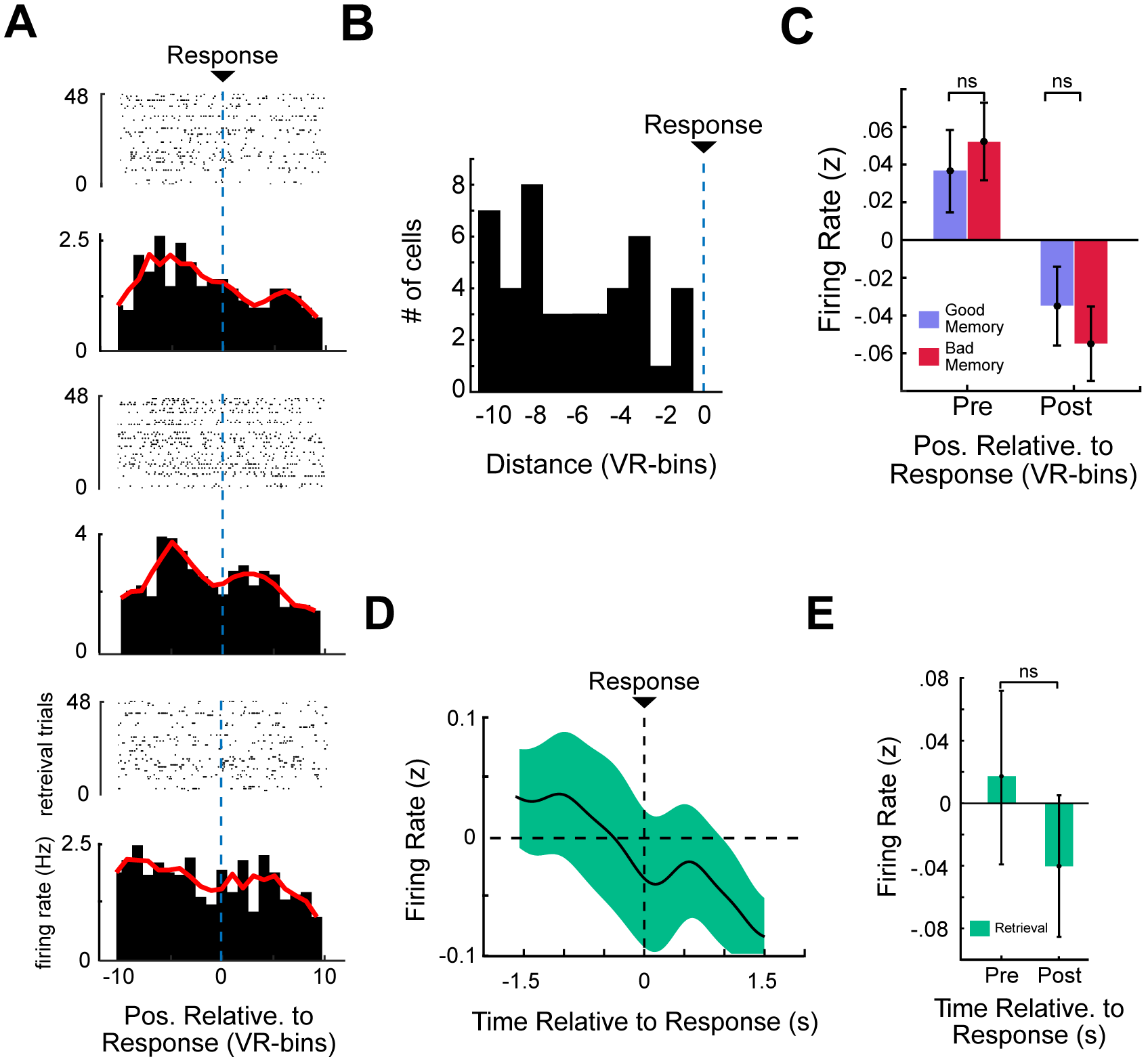
Trace-cell activity increases relative to response location, and not time. A) Raster plot and corresponding PSTH of three representative trace cells, aligned relative to response location (indicated by blue dotted line). B) Mean firing rate (z-scored) of all trace cells aligned to response time. In contrast to Fig. 2F, firing rate here is assessed as a function of time surrounding button press, rather than spatial bin. Notably, these cells do not show significant response period activity at consistent times preceding button press, implying that trace cells were not simply activating at fixed times preceding the button press (t-test by spatial bin, *p >* 0.05, FDR-corrected C) Pre- and post-response trace-cell firing rate, binned by time as in Panel *B*. D) Mean firing rate (z-scored) of all trace cells aligned to response time. Shading indicates SEM. No bins show significant difference from baseline. E) Pre-response and post-response firing rate (z-scored) relative to response time. ”Ns” indicate non-significance from a t-test (*p >* 0.05).

**Supplementary Figure 4:**
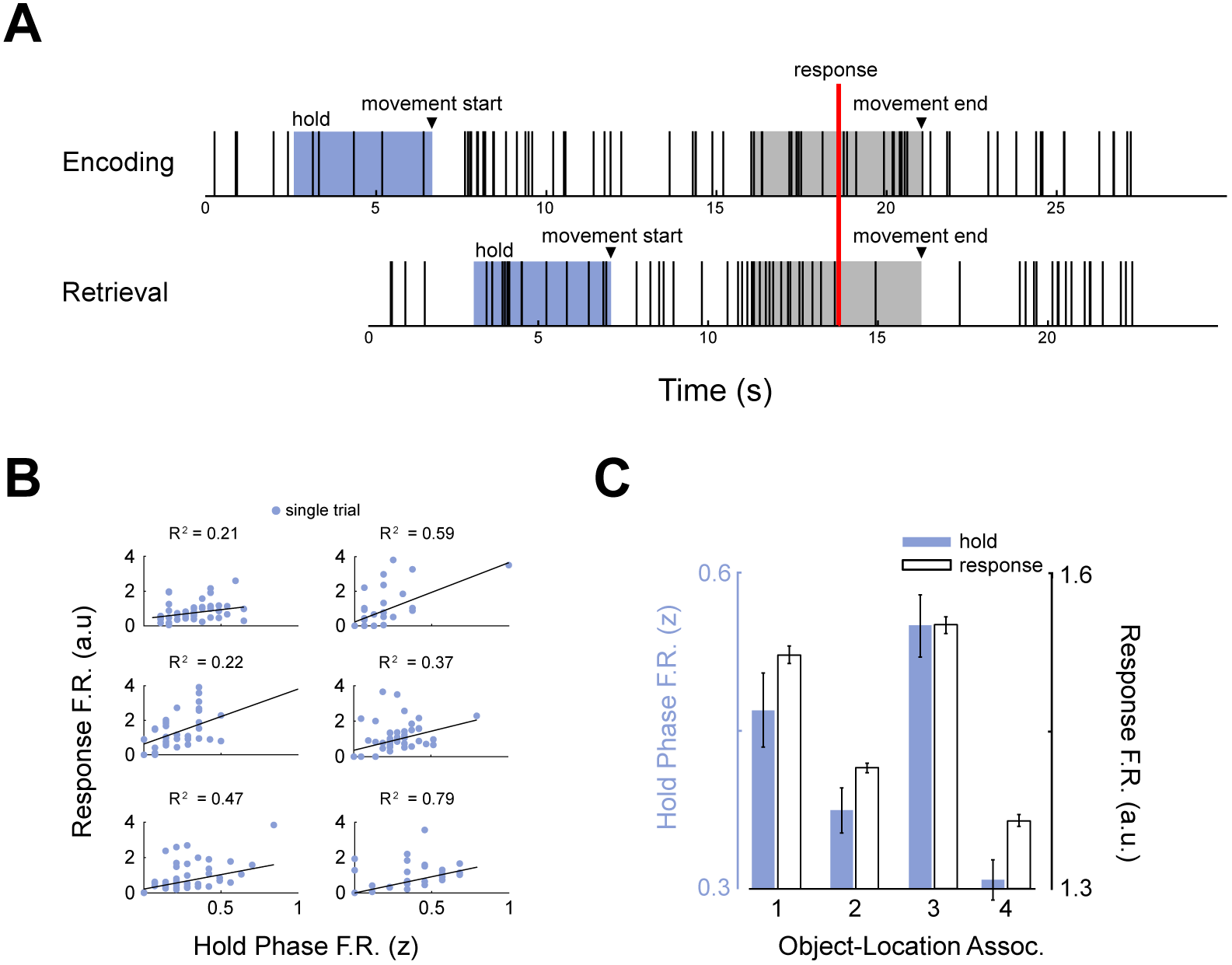
Examples of trace cell activity during hold period and response period: A) Raster plot indicating the activity of an entorhinal trace cell during encoding (top) and retrieval (bottom) trials for a particular object cue. Vertical lines denote spike times. Blue shading indicates the hold period. Arrows denote the start and end of the movement period. Grey shading indicates 5 s around button press. Note that this cell shows an increase in firing rate during the hold period. Increases in activity are also visible during the response period preceding response, with the cell largely ceasing to fire following the response. These encoding and retrieval trials have different durations, which is a result from the differing movement speeds on the two trials. B) Scatter plots illustrating the relations between normalized firing rates in the hold and response periods for six representative trace cells. Black line denotes the robust linear regression fit. C) Hold period firing rate and response period firing rate, for a representative EC trace cell, averaged across all trials for each cue object.

**Supplementary Figure 5:**
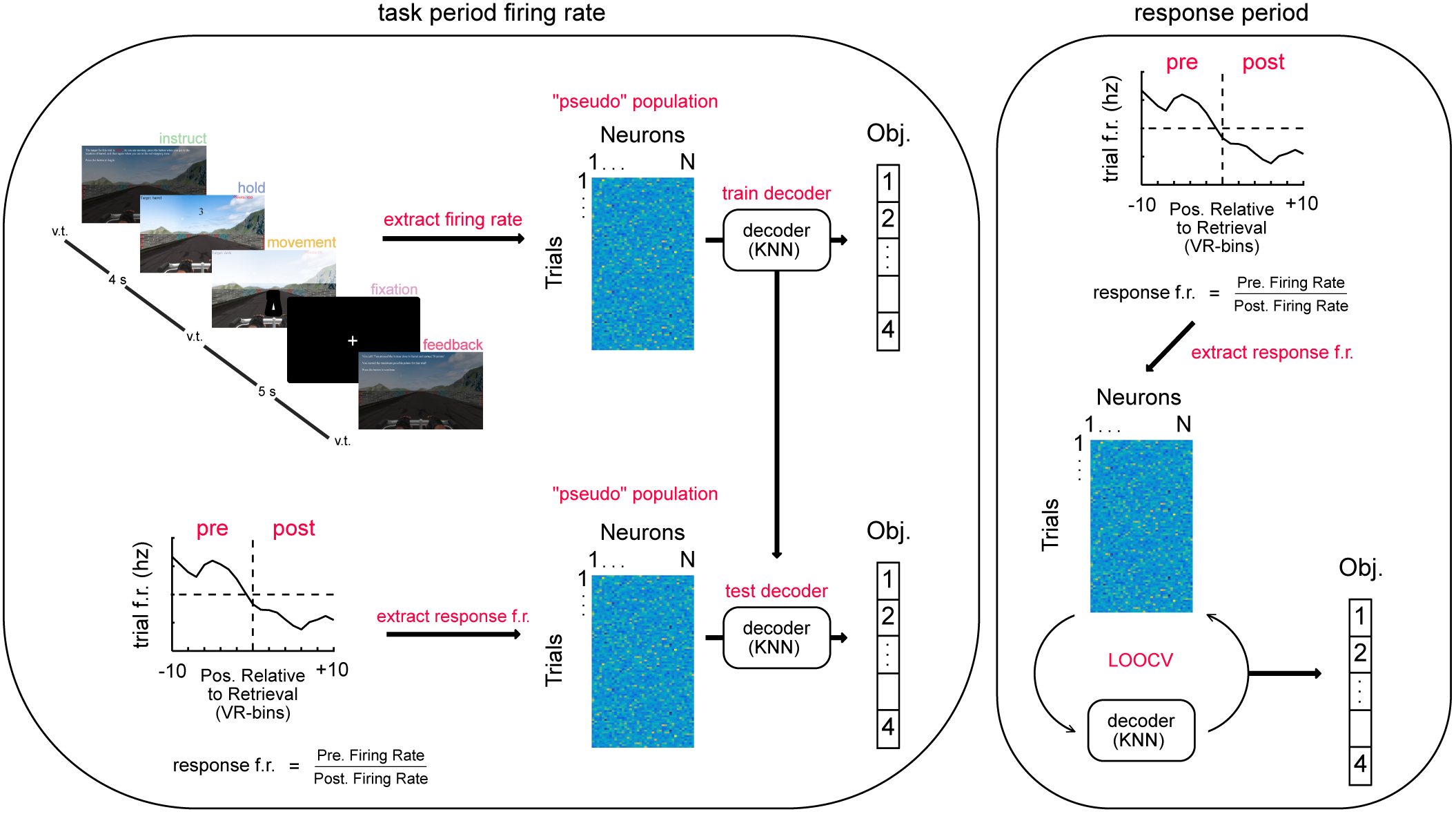
Illustration of the multivariate decoding procedure. Left, top: **Training:** Pseudopopulations of neural activity were constructed by extracting the activity of cells during the different period of interest. Decoders were trained on this data using a *k* -nearest-neighbor (KNN) framework to predict the object cue for each trial. Left, bottom: **Testing:** Response period activity for each trial was computed by normalizing the pre-response firing rate by the post-response firing rate. This measure was extracted for each trial and used as the test set for the decoders trained on each task period. Right: **LOOCV decoder using response period.** We also trained and tested a decoder using just the response period activity. In order to ensure we had separate train–test data, we used leave-one-out cross-validation (LOOCV). Feature extraction and decoding framework were consistent with those used for the task periods(Left).

**Supplementary Table 1:** Contribution of subjects and sessions to total cell counts: Table indicates the total number of trace cells and total cells for each patient, by brain region. The right-most column indicates the number of trace cells observed on unique recording channels across sessions of the task

| Subject # | # Sessions | # Amygdala units | # Hippocampal units | # Entorhinal units | # Cingulate units | # Trace cells |
| --- | --- | --- | --- | --- | --- | --- |
| R1008J | 1 | 0 of (6) | 0 of (6) | 0 of (0) | 0 of (0) | 0 |
| R1027J | 1 | 1 of (15) | 0 of (1) | 0 of (0) | 0 of (0) | 1 |
| R1030J | 4 | 1 of (21) | 0 of (1) | 0 of (0) | 0 of (0) | 1 |
| R1092J | 3 | 0 of (0) | 0 of (17) | 0 of (0) | 0 of (0) | 0 |
| R1139C | 2 | 0 of (0) | 0 of (0) | 0 of (0) | 2 of (10) | 2 |
| R1152C | 1 | 0 of (0) | 0 of (2) | 0 of (0) | 5 of (14) | 5 |
| R1182C | 1 | 0 of (0) | 0 of (0) | 1 of (4) | 1 of (3) | 2 |
| R1219C | 1 | 0 of (0) | 0 of (6) | 3 of (14) | 0 of (0) | 3 |
| R1278E | 3 | 0 of (0) | 0 of (0) | 20 of (65) | 0 of (0) | 20 |
| UT048 | 1 | 0 of (10) | 1 of (2) | 0 of (0) | 0 of (0) | 1 |
| R1268T | 1 | 0 of (0) | 0 of (2) | 0 of (0) | 0 of (0) | 0 |
| R1241J | 1 | 1 of (10) | 0 of (0) | 0 of (0) | 0 of (0) | 1 |
| R1297T | 1 | 0 of (0) | 0 of (3) | 0 of (0) | 0 of (0) | 0 |
| R1299T | 1 | 0 of (0) | 1 of (12) | 0 of (0) | 0 of (0) | 1 |
| R1315T | 2 | 0 of (0) | 0 of (12) | 0 of (0) | 0 of (0) | 0 |
| EU001 | 1 | 0 of (0) | 1 of (19) | 0 of (0) | 0 of (0) | 1 |
| R1354E | 2 | 0 of (0) | 2 of (14) | 0 of (0) | 0 of (0) | 2 |
| R1362E | 3 | 0 of (0) | 0 of (0) | 4 of (26) | 0 of (0) | 4 |
| R1414E | 1 | 0 of (0) | 0 of (0) | 1 of (4) | 0 of (0) | 1 |

